# The devil is in the detail: The role of threat level and intolerance of uncertainty in extinction

**DOI:** 10.1101/479543

**Authors:** Jayne Morriss, Francesco Saldarini, Carien M. van Reekum

## Abstract

Recent evidence suggests that individual differences in intolerance of uncertainty (IU) are associated with disrupted threat extinction. However, it is unknown what maintains the learned threat association in high IU individuals: is it the experienced uncertainty during extinction or the combination of experienced uncertainty with potential threat during extinction? Here we addressed this question by running two independent experiments with uncertain auditory stimuli that varied in threat level (Experiment 1, aversive human scream (*n* = 30); Experiment 2, benign tone (*n* = 47) and mildly aversive tone (*n* = 49)). During the experiments, we recorded skin conductance responses and subjective ratings to the learned cues during acquisition and extinction. In experiment 1, high IU was associated with heightened skin conductance responding to the learned threat vs. safe cue during extinction. In experiment 2, high IU was associated only with larger skin conductance responding to the learned cues with threatening properties during extinction i.e. mildly aversive tone. These findings suggest that uncertainty in combination with threat, even when mild, disrupts extinction in high IU individuals. Such findings help us understand the link between IU and threat extinction, and its relevance to anxiety disorder pathology.

## Introduction

Adjusting behaviour based on predictive cues that signal threat and safety is adaptive (LeDoux & Daw, 2018). An organism can learn to associate neutral cues (conditioned stimulus, e.g. a visual stimulus such as a shape) with threatening outcomes (unconditioned stimulus, e.g. shock, loud tone) or safe outcomes. Repeated presentations of a neutral cue with a threatening outcome results in defensive responding to the conditioned cue. This learned association can also be extinguished by repeatedly presenting the conditioned cue without the aversive outcome, resulting in a reduction in defensive responding. Partial reinforcement of aversive stimuli (e.g. shock, noise), particularly at 50% reinforcement rate, has been shown to maintain the conditioned response during extinction (Leonard, 1975; Livneh & Paz, 2012). This effect is known as the partial reinforcement extinction effect (PREE). After partial reinforcement, it is thought that the conditioned response is maintained during extinction due to the uncertainty of receiving a threatening outcome (for discussion see Li, Ishii & Naoki, 2016).

Overestimating the predictability of threat over safety is a common feature of anxiety and stress disorders (Duits et al., 2015; Milad & Quirk, 2012). A large body of research has shown that individuals who have anxious traits or who are clinically anxious show reduced extinction of threat, indexed by larger physiological responses to cues that no longer predict an aversive outcome (Etkin & Wager, 2007; Lonsdorf & Merz, 2017). Emerging research from our lab and others suggest that individual differences in intolerance of uncertainty (IU), the tendency to find uncertainty anxiety provoking, may play a specific role in maintaining threat bias during extinction (Dunsmoor, Campese, Ceceli, LeDoux, & Phelps, 2015; Lucas, Luck, & Lipp, 2018; Morriss, Christakou, & van Reekum, 2015, 2016; Morriss, Macdonald, & van Reekum, 2016). For example, after 100% reinforcement, high IU, relative to low IU individuals have been found to show generalized skin conductance response across threat and safety cues during early extinction, and to show continued skin conductance responding to threat versus safety cues during late extinction (Morriss, Christakou, & van Reekum, 2015, 2016). Moreover, after 50% reinforcement, high IU has been found to be associated with generalized skin conductance responding to parametrically graded stimuli during extinction (e.g. stimuli that vary in similarity to the learned threat cue) (Morriss, Macdonald, & van Reekum, 2016). Individual differences in IU are typically associated with responding during the extinction phase and not during the acquisition phase (but see Chin et al., 2016; Morriss, Macdonald, & van Reekum, 2016).

During extinction, there is a period of uncertainty regarding the change of outcome i.e. threat to safe, and this may induce anxiety in high IU individuals. The extent of uncertainty-induced anxiety experienced by high IU individuals during extinction may vary as a function of threat level; in the case of extinction, this could be the level of aversiveness of the US. However, it is unknown as to: (1) whether high IU individuals would exhibit disrupted extinction in the absence of threat, as uncertainty is aversive enough in itself, or (2) whether some type of threat, even when mild, is required. This question can be examined by varying the level of threat during extinction in independent samples. Given the important role of uncertainty in anxiety (Carleton, 2016a, 2016b; Grupe & Nitschke, 2013) and that current psychological therapies are based on associative learning principles (Craske, Treanor, Conway, Zbozinek, & Vervliet, 2014; Milad & Quirk, 2012), examining the parameters by which extinction leads to uncertainty-induced anxiety in high IU individuals may provide crucial information relevant to anxiety disorder pathology and treatment.

We conducted two experiments using threat and safety cues during acquisition and extinction. For each experiment, we varied the properties of the unconditioned stimulus to assess the relationship between individual differences in self-reported IU and the level of threat during extinction. In the first experiment, we aimed to replicate previous IU and extinction findings using an aversive human scream as the unconditioned stimulus with a 50% reinforcement schedule (Morriss, Christakou, & van Reekum, 2015, 2016; Morriss, Macdonald, & van Reekum, 2016). In the second experiment, we aimed to examine the extent to which IU would predict reduced extinction when using different unconditioned stimuli that varied in aversiveness i.e. mildly aversive to neutral tones. In experiment 2, we tested two independent samples of participants, with each being presented one of the tones. During both experiments, we measured skin conductance response (SCR) and expectancy ratings whilst participants performed the acquisition and extinction phases. We used sounds as unconditioned stimuli and visual shape stimuli as conditioned stimuli, similar to previous conditioning research (Neumann, Waters, & Westbury, 2008; Phelps, Delgado, Nearing, & LeDoux, 2004). We used a 50% reinforcement rate during acquisition to maintain conditioning (Leonard, 1975; Livneh & Paz, 2012) and induce greater uncertainty during extinction (Li, Ishii & Naoki, 2016), similar to our previous work (Morriss, Macdonald, & van Reekum, 2016).

In general for experiments 1 and 2, we hypothesised that there would be greater skin conductance responding and expectancy ratings to the learned uncertain (threat, mild threat, neutral, also known as the CS+) versus certain (safe, also known as the CS-) cues during acquisition. In addition, for experiment 1, we hypothesised that high IU would be associated with (1) greater skin conductance responding to both the CS+ and CS-cues during early extinction, and (2) greater skin conductance responding to the CS+ versus CS-during late extinction (Morriss, Christakou, & van Reekum, 2015, 2016), suggesting compromised updating of the CS+ to safe in individuals reporting high IU. For experiment 2, we had two exploratory hypotheses for IU and updating of learned associations during extinction: (1) If uncertainty is aversive enough in itself, we expected high IU, relative to low IU, to predict greater skin conductance responding to the CS+ versus the CS-, regardless of aversiveness of the unconditioned stimulus. (2) If some level of threat is required, we expected high IU, relative to low IU to only predict greater skin conductance responding to the CS+ with mild threat versus the CS+ signalling a more neutral outcome. For both acquisition and extinction, we tested the specificity of IU effects by controlling for individual variation reported on the commonly used Spielberger State-Trait Anxiety Inventory, Trait Version (STAI) (Spielberger, Gorsuch, Lushene, Vagg, & Jacobs, 1983). We did not have specific predictions for individual differences in STAI or IU predicting expectancy ratings, as previous experiments in our lab have not found consistent results for expectancy ratings (Morriss, Christakou, & van Reekum, 2016; Morriss, MacDonald, & van Reekum, 2016).

## Experiment 1: Method

### Participants

Thirty volunteers (*M* age = 23.53, *SD* age = 4.96; 16 females and 14 males) took part in the study. All participants had normal or corrected to normal vision.

Participants provided written informed consent and received £5 for their participation. Advertisements and word of mouth were used to recruit participants from the University of Reading and local area. The procedure was approved by the University of Reading Research Ethics Committee.

### Procedure

Participants completed questionnaires online before the study. On the day of the experiment participants arrived at the laboratory and were informed on the experimental procedures. Firstly, participants were seated in the testing booth and asked to complete a consent form as an agreement to take part in the study. Secondly, physiological sensors were attached to the participants’ non-dominant hand. The conditioning task (see “Conditioning task” below for details) was presented on a computer, whilst skin conductance, interbeat interval and behavioural ratings were recorded. Participants were instructed to: (1) maintain attention to the task by looking at the coloured squares and listening to the sounds, which may be unpleasant, (2) respond to the expectancy rating scales that followed each block of trials, using number keys on the keyboard with their dominant hand and (3) to stay as still as possible. The experiment took approximately 30 minutes in total.

### Conditioning task

The conditioning task was designed using E-Prime 2.0 software (Psychology Software Tools Ltd, Pittsburgh, PA). Visual stimuli were presented at a 60 Hz refresh rate with a 800 × 600 pixel resolution. Participants sat approximately 60 cm from the screen. Visual stimuli were blue and yellow squares with 183 × 183 pixel dimensions that resulted in a visual angle of 5.78° × 9.73°. The aversive sound stimulus was presented through headphones. The sound consisted of a fear inducing female scream (for sound parameters, see Morriss, Christakou & van Reekum, 2015). The volume of the sound was standardized across participants by using fixed volume settings on the presentation computer and was verified by an audiometer prior to each session.

The task comprised of two learning phases: acquisition and extinction. Both acquisition and extinction consisted of two blocks. In acquisition, one of the coloured squares (blue or yellow) was paired with the aversive 90 dB sound 50% of the time (CS+), whilst the other square (yellow or blue) was presented alone (CS-). The 50% pairing rate was designed to maximize uncertainty of the CS+ / US contingency. During extinction, both the blue and yellow squares were presented in the absence of the US.

The acquisition phase consisted of 24 trials (6 CS+ paired, 6 CS+ unpaired, 12 CS-) and the extinction phase 32 trials (16 CS+ unpaired, 16 CS-). Early extinction was defined at the first 8 CS+/CS-trials and late extinction was defined as the last 8 CS+/CS-trials. Experimental trials were pseudo-randomized such that the first trial of acquisition was always paired and then after all trial types were randomly presented. Conditioning contingencies were counterbalanced, with half of participants receiving the blue square paired with the US and the other half of participants receiving the yellow square paired with the US. The coloured squares were presented for a total of 4000 ms. The aversive sound lasted for 1000 ms, which coterminated with the reinforced CS+’s. Subsequently, a blank screen was presented for 6000 – 8800 ms.

At the end of each block (4 blocks in total, 2 in acquisition and 2 in extinction), participants were asked to rate how much they expected the blue square and yellow square to be followed by the sound stimulus, where the scale ranged from 1 (“Don’t Expect”) to 9 (“Do Expect”). Two other 9-point Likert scales were presented at the end of the experiment. Participants were asked to rate the valence and arousal of the sound stimulus. The scales ranged from 1 (Valence: very negative; Arousal: calm) to 9 (Valence: very positive; Arousal: excited).

### Questionnaires

To assess anxious disposition, we administered the Spielberger State-Trait Anxiety Inventory – Trait version (STAI) and Intolerance of Uncertainty (IU) questionnaire (Freeston, Rhéaume, Letarte, Dugas, & Ladouceur, 1994). The IU measure consists of 27 items, example items include “Uncertainty makes me uneasy, anxious, or stressed” and “I must get away from all uncertain situations”. Similar distributions and internal reliability of scores were found for the anxiety measures, STAI (*M* = 41.30; *SD* = 9.84; range = 26-56; α = .91), IU (*M* = 67.50; *SD* = 17.18; range = 33-94; α = .93).

### Behavioural data scoring

Rating data were reduced for each participant by calculating their average responses for each experimental condition (Acquisition CS+; Acquisition CS-; Extinction CS+ Early; Extinction CS-Early; Extinction CS+ Late; Extinction CS-Late) using the E-Data Aid tool in E-Prime (Psychology Software Tools Ltd, Pittsburgh, PA).

### Physiological acquisition and scoring

Physiological recordings were obtained using AD Instruments (AD Instruments Ltd, Chalgrove, Oxfordshire) hardware and software. Electrodermal activity was measured with dry MLT116F silver/silver chloride bipolar finger electrodes that were attached to the distal phalanges of the index and middle fingers of the non-dominant hand. A low constant-voltage AC excitation of 22 mV_rms_ at 75 Hz was passed through the electrodes, which were connected to a ML116 GSR Amp, and converted to DC before being digitized and stored. Interbeat Interval (IBI) was measured using a MLT1010 Electric Pulse Transducer, which was connected to the participant’s distal phalange of the ring finger. An ML138 Bio Amp connected to an ML870 PowerLab Unit Model 8/30 amplified the skin conductance and IBI signals, which were digitized through a 16-bit A/D converter at 1000 Hz. IBI signal was used only to identify movement artefacts and was not analysed. The electrodermal signal was converted from volts to microSiemens using AD Instruments software (AD Instruments Ltd, Chalgrove, Oxfordshire).

CS+ unpaired and CS-trials were included in the analysis, but CS+ paired trials were discarded to avoid sound confounds. Skin conductance responses (SCR) were scored when there was an increase of skin conductance level exceeding 0.03 microSiemens. The amplitude of each response was scored as the difference between the onset and the maximum deflection prior to the signal flattening out or decreasing. SCR onsets and respective peaks were counted if the SCR onset was within 0.5-3.5 seconds following CS onset. Trials with no discernible SCRs were scored as zero (Morriss, Chapman, Tomlinson, & van Reekum, 2018). SCR’s were square root transformed to reduce skew at the trial level (Dawson, Schell, & Filion, 2000) and were z-scored to control for interindividual differences in skin conductance responsiveness (Ben-Shakhar, 1985). SCR magnitudes were calculated by averaging the transformed values for each condition, creating the following conditions: Acquisition CS+; Acquisition CS-; Extinction CS+ Early; Extinction CS-Early; Extinction CS+ Late; Extinction CS-Late.

### SCR magnitude inclusion

In the sample, we had one non-responder, defined as having less than 10% of SCR responses to unpaired trials across acquisition and extinction. We report below the SCR magnitude results without the non-responder included.

### Ratings and SCR magnitude analysis

The analysis was conducted using the mixed procedure in SPSS 21.0 (SPSS, Inc; Chicago, Illinois). We conducted separate multilevel models on ratings and SCR magnitude for each phase (Acquisition, Extinction). For ratings and SCR magnitude during the acquisition phase we entered Stimulus (CS+, CS-) at level 1 and individual subjects at level 2. For ratings and SCR magnitude during the extinction phase we entered Stimulus (CS+, CS-) and Time (Early, Late) at level 1 and individual subjects at level 2. We included the following individual difference predictor variables into the multilevel models: IU and STAI. In all models, we used a diagonal covariance matrix for level 1. Random effects included a random intercept for each individual subject, where a variance components covariance structure was used. Fixed effects included Stimulus, Phase and Time. We used a maximum likelihood estimator for the multilevel models. We used the least significance difference procedure for pairwise comparisons.

In the model where there are two predictor variables (IU, STAI), a significant interaction with one variable but not the other suggests specificity. Based on our prior work, we expected such specificity for IU, but we explored interactions with STAI, given extant findings with STAI in the conditioning literature (e.g. Lonsdorf & Merz, 2017). Where a significant interaction was observed with IU (or STAI), we performed follow-up pairwise comparisons on the estimated marginal means of the relevant conditions estimated at specific IU values of + or −1 SD of mean IU, adjusted for STAI (or IU). These data are estimated from the multilevel model of the entire sample, not unlike performing a simple slopes analysis in a multiple regression analysis. Similar analyses have been published elsewhere (Morriss, Macdonald, & van Reekum, 2016; Morriss, McSorley, & van Reekum, 2017).

### Experiment 1: Results

For descriptive statistics see Table 1.

**Table 1.** Experiment 1 summary of means (SD) for each dependent measure as a function of condition (CS+ and CS-), separately for acquisition, early extinction and late extinction.

| Measure | Acquisition |  | Early Extinction |  | Late Extinction |  |
| --- | --- | --- | --- | --- | --- | --- |
|  | CS+ | CS- | CS+ | CS- | CS+ | CS- |
| Square root transformed and z-scored SCR magnitude ($\sqrt{\mu\text{S}}$) | 0.22<br>(0.30) | 0.04<br>(0.24) | 0.10<br>(0.44) | -0.16<br>(0.41) | -0.05<br>(0.31) | -0.11<br>(0.43) |
| Expectancy rating (1-9) | 4.43<br>(1.26) | 3.33<br>(1.62) | 6.77<br>(1.22) | 2.93<br>(1.89) | 1.90<br>(1.65) | 1.90<br>(1.92) |

Note: SCR magnitude ( $\sqrt{\mu\text{S}}$ ), square root transformed and z-scored skin conductance magnitude measured in microSiemens.

### Ratings

Participants rated the human scream sound stimulus as aversive (*M* = 2.43 *SD* = 1.41, range 1-7, where 1 = very negative and 9 = very positive) and arousing (*M* = 6.50, *SD* = 1.78, range 2-9 where 1 = calm and 9 = excited).

Participants had higher expectancy ratings of the sound with the CS+ versus CS-during acquisition [Stimulus, *F*(1, 30) = 16.075, *p* < .001] and extinction [Stimulus, *F*(1, 81.159) = 65.290, *p* < .001; see Table 1]. In the early part of extinction, participants displayed higher expectancy ratings of the sound with the CS+ versus CS-, *p* = .001. However, during late extinction, the expectancy rating of the sound with the CS+ dropped and was similar to the CS-, *p* > .5 [Time, *F*(1, 81.159) = 154.667, *p* < .001; Stimulus × Time, *F*(1, 81.159) = 65.290, *p* < .001].

During acquisition, individuals scoring lower in STAI tended to have greater discrimination between expectancy of the sound with the CS+ (*M* = 5.05, *SE* = .41) versus CS-(*M* = 2.70, *SE* = .54), *p* < .001, whilst individuals higher STAI tended to have poorer discrimination between expectancy of the sound with the CS+ (*M* = 3.82, *SE* = .41) and CS-(*M* = 3.96, *SE* = .54), *p* = .781 [Stimulus × STAI, *F*(1, 30) = 4.141, *p* = .026]. Moreover, during extinction, the same pattern of discrimination was observed, as low STAI showed greater discrimination, *p* < .001 (CS+: *M* = 4.32, *SE* = .44; CS-: *M* = 1.58, *SE* = .46), compared to high STAI. *p* = .02 (CS+: *M* = 4.33, *SE* = .44; CS-: *M* = 3.25, *SE* = .46) [Stimulus × STAI, *F*(1, 81.159) = 4.493, *p* = .037]. A similar pattern was observed for IU during extinction but was not statistically as strong, [Stimulus × IU, *F*(1, 81.159) = 4.146, *p* = .045]. No other significant main effects or interactions with STAI or IU were found, max *F* = 3.780.

### SCR magnitude

Larger average SCR magnitude was found for the CS+, compared to the CS-during acquisition [Stimulus, *F*(1, 29) = 9.029, *p* = .005]. Unexpectedly, during acquisition, high STAI was associated with greater SCR magnitude to CS+ (*M* = .256, *SE* = .105) vs. CS-(*M* = -.137, *SE* = 071), *p* = .003, whilst low STAI was associated with reduced SCR magnitude difference between CS+ (*M* = .186, *SE* = .097) vs. CS (*M* = .193, *SE* = .066), *p* = .914 [Stimulus × STAI, *F*(1, 29) = 4.294, *p* = .023]^1^. Individual differences in IU were not associated with SCR during this phase.

During extinction, larger SCR magnitude was observed for the CS+ versus CS-[Stimulus, *F*(1, 108.786) = 5.736, *p* = .018; see Table 1]. Partially in line with our predictions, higher IU was associated with greater SCR magnitude response to the CS+ versus CS-during extinction, *p* < .001, whilst lower IU was associated with no significant differential SCR magnitude response between the CS+ and CS-, *p* = .239 [Stimulus × IU, *F*(1, 108.786) = 8.351, *p* = .005] (see Figure 1). Time (early vs late) did not affect this relationship, however. No other significant main effects or interactions with IU or STAI were found, max *F* = 2.279.

**Fig 1.**
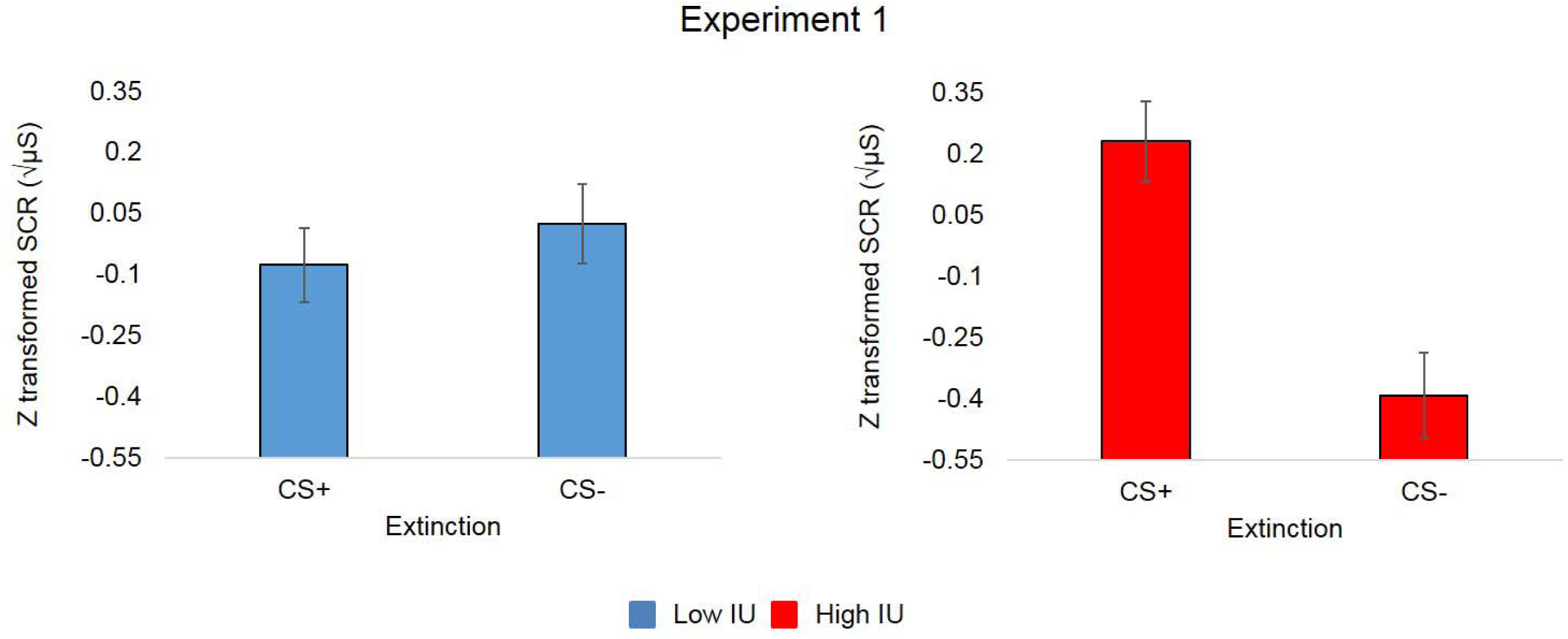
Bar graphs depicting IU estimated at + or - 1 SD of mean IU (controlling for STAI) from the multilevel model analysis for SCR magnitude during extinction. In experiment 1, high IU, relative to low IU individuals were found to show heightened skin conductance responding to the CS+ versus CS-cue during extinction. Bars represent standard error at + or – 1 SD of mean IU. Square root transformed and z-scored SCR magnitude (μS), skin conductance magnitude measured in microSiemens.

## Experiment 1: Conclusion

For experiment 1 we observed typical profiles of acquisition and extinction, where larger SCR magnitudes and expectancy ratings were found for the CS+ vs. CS-.

High IU was associated with larger SCR magnitude to the CS+ vs. CS-during extinction. This finding partially replicates our previous research (Morriss, Christakou, & Van Reekum, 2015, 2016), as we did not observe time-based effects of IU and threat extinction. In our original experiments of IU and threat extinction, we observed high IU individuals to display generalized skin conductance response across CS+ and CS-cues during early extinction, and to show continued skin conductance responding to CS+ versus CS-cues during late extinction (Morriss, Christakou, & van Reekum, 2015, 2016).

In addition, we observed greater discrimination of expectancy ratings of the sound with the CS+ vs. CS-during acquisition and extinction for individuals lower in STAI. However, during acquisition individuals lower in STAI showed reduced discrimination in SCR magnitude for the CS+ vs. CS-.

## Experiment 2: Method

All aspects of the method are identical to experiment 1, except the following below.

### Participants

Ninety six volunteers (*M* age = 19.59, *SD* age = 1.93; 81 females and 15 males) took part in the study. The neutral tone group, N =47 (M age = 19.28, SD age = 1.16; 38 females and 9 males), and aversive tone group, N = 49 (M age = 19.89, SD age = 2.43; 43 females and 6 males) underwent similar conditioning procedures, but received different US stimulation (see “Conditioning task” below for details). All participants had normal or corrected to normal vision. Participants provided written informed consent and received 0.5 credits for their participation. The procedure was approved by the University of Reading Research Ethics Committee.

### Procedure

On the day of the experiment participants arrived at the laboratory and were informed on the experimental procedures. Firstly, participants were seated in the testing booth and asked to complete a consent form as an agreement to take part in the study and a set of questionnaires on the computer (see below). Group allocation was based on the average IU score from previous studies in our lab (Morriss, Christakou, & van Reekum, 2016; Morriss & van Reekum, unpublished). Participants with low (below average < 65) and high IU (above average > 65) were evenly distributed to the neutral tone and aversive tone groups. Next, physiological sensors were attached to the participants’ non-dominant hand. The conditioning task (see “Conditioning task” below for details) was presented on a computer, whilst skin conductance, interbeat interval (to help in artefact detection in skin conductance) and behavioural ratings were recorded. Participants were instructed to: (1) maintain attention to the task by looking at the coloured squares and listening to the sounds, which may be unpleasant, (2) respond to the expectancy rating scales that followed each block of trials, using number keys on the keyboard with their dominant hand and (3) to stay as still as possible. The experiment took approximately 30 minutes in total.

### Conditioning task

The conditioning task procedure in experiment 2 was similar to experiment 1. Visual stimuli were blue and yellow squares presented on a computer screen and served as CSs (see “Experiment 1: Method”, “Conditioning task” section for more details). The aversive sound stimulus was presented through headphones and served as US. Each experimental group received a different auditory stimulus. The Aversive Tone Group was exposed to a high pitched tone (1600 Hz, 1000 ms, 90 db). The Neutral Tone Group was exposed to a low pitched tone (360 Hz, 1000 ms, 80 db). We used Audacity 2.0.3 software (http://audacity.sourceforge.net/) to generate the tones. The volume of the sound was standardized across participants by using fixed volume settings on the presentation computer and was verified by an audiometer prior to each session.

### Questionnaires

Similar distributions and internal reliability of scores were found for the anxiety measures. For the neutral tone group: STAI (*M* = 45.40; *SD* = 9.78; range = 29-66; α = .91), IU (*M* = 68.21; *SD* = 15.04; range = 42-101; α = .90). For the aversive tone group: STAI (*M* = 42.55; *SD* = 10.89; range = 26-70; α = .92), IU (*M* = 67.94; *SD* = 15.59; range = 42-110; α = .91). The groups did not significantly differ on STAI or IU scores [*t’s* < 1.4, *p’*s > .15].

### SCR magnitude inclusion

Based on the criterion specified in experiment 1, the neutral tone group had seven non-responders, and the aversive tone group had ten non-responders. This left forty participants in the neutral tone group and thirty-nine participants in the aversive tone group with usable SCR data. We report below the SCR magnitude results excluding the non-responders.

### Ratings and SCR magnitude analysis

The same statistical procedures from Experiment 1 were used in Experiment 2. We added an additional factor of Group (Neutral tone, Aversive tone).

## Experiment 2: Results

For descriptive statistics see Table 2.

**Table 2.** Experiment 2 summary of means (SD) for each dependent measure as a function of group (Neutral Tone and Aversive Tone) and condition (CS+ and CS-), separately for acquisition, early extinction and late extinction.

| Group | Measure | Acquisition |  | Early Extinction |  | Late Extinction |  |
| --- | --- | --- | --- | --- | --- | --- | --- |
|  |  | CS+ | CS- | CS+ | CS- | CS+ | CS- |
| Neutral Tone | Square root transformed and z-scored SCR magnitude ($\sqrt{\mu\text{S}}$) | 0.18 | 0.01 | 0.2 | -0.11 | -0.01 | -0.04 |
|  |  | (0.43) | (0.28) | (0.38) | (0.28) | (0.44) | (0.39) |
|  | Expectancy rating (1-9) | 5.86 | 1.84 | 3.28 | 1.66 | 2.11 | 1.49 |
|  |  | (1.49) | (1.39) | (2.00) | (1.59) | (1.49) | (1.23) |
| Aversive Tone | Square root transformed and z-scored SCR magnitude ($\sqrt{\mu\text{S}}$) | 0.17 | 0.06 | 0.01 | -0.02 | -0.04 | -0.18 |
|  |  | (0.57) | (0.29) | (0.29) | (0.31) | (0.40) | (0.30) |
|  | Expectancy rating (1-9) | 5.99 | 1.84 | 3.20 | 1.76 | 2.10 | 1.35 |
|  |  | (1.55) | (1.53) | (1.72) | (1.85) | (1.58) | (1.30) |

Note: SCR magnitude ( $\sqrt{\mu\text{S}}$ ), square root transformed and z-scored skin conductance magnitude measured in microSiemens.

### Ratings

In the neutral tone group, the sound was rated as slightly aversive (*M* = 4.08, *SD* = 1.28, range 2-8, where 1 = very negative and 9 = very positive) and neutral in arousal (*M* = 5.02, *SD* = 1.76, range 1-8 where 1 = calm and 9 = excited). In the aversive tone group, the sound was rated as moderately aversive (*M* = 3.42 *SD* = 1.45, range 1-7, where 1 = very negative and 9 = very positive) and arousing (*M* = 5.97, *SD* = 1.78, range 1-8 where 1 = calm and 9 = excited). The aversive tone was rated significantly more aversive [*t =* −2.636, *p* = .010] and arousing [*t =* 2.339, *p* = .021] than the neutral tone.

Participants had higher expectancy ratings of the tones with the CS+ versus CS-during acquisition [Stimulus, *F*(1, 175.914) = 339.935, *p* < .001; see Table 2]. No other significant effects of Group or interactions with Group, IU or STAI were observed during acquisition, Max *F* = 1.021. Similar patterns of ratings during extinction were observed for the neutral and aversive tone groups [Stimulus, *F*(1, 240.054) = 94.134, *p* < .001; Time, *F*(1, 240.054) = 40.569, *p* < .001; Stimulus × Time, *F*(1, 240.054) = 13.329, *p* < .001]. Participants displayed higher expectancy ratings of the tones with the CS+ versus CS-during early extinction, compared to late extinction, *p’*s <.001.

Surprisingly, during extinction, an effect of STAI was found [Stimulus × STAI, *F*(1, 240.054) = 3.961, *p =* .048], where low STAI was associated with greater discrimination of expectancy of the tones with the CS+ (*M* = 2.98, *SE* = .26) vs. CS-(*M* = 1.54, *SE* = .24), *p* < .001, compared to high STAI, *p* <.001 (CS+: *M* = 2.42, *SE* = .26; CS-: *M* = 1.62, *SE* = .24). No other significant main effects or interactions were found during extinction, max *F* = 2.907.

### SCR magnitude

Greater SCR magnitude was found for the CS+, compared to the CS-during acquisition [Stimulus, *F*(1, 71) = 5.079, *p* = .027; see Table 2]. No other significant main effects of Group or interactions with IU or STAI emerged during acquisition, max *F* = 1.301.

Larger SCR magnitude was found for the CS+, compared to the CS-during extinction [Stimulus, *F*(1, 287.741) = 4.924, *p* = .027]. Notably, tone group and individual differences in IU predicted SCR magnitude during extinction [Stimulus × Group × IU, *F*(1, 287.741) = 5.279, *p* = .022]. In the aversive tone group, higher IU was associated with greater SCR magnitude response during extinction to the CS+ versus CS-, *p* = .009, whilst lower IU was associated with no significant differential SCR magnitude response during extinction between the CS+ and CS-, *p* = .489 (see Figure 2). Interestingly, in the neutral tone group, higher IU was associated with no significant differential SCR magnitude response during extinction to the CS+ versus CS-, *p* = .864, whilst lower IU was associated with a significant differential SCR magnitude response during extinction between the CS+ and CS-, *p* = .041.

**Fig 2.**
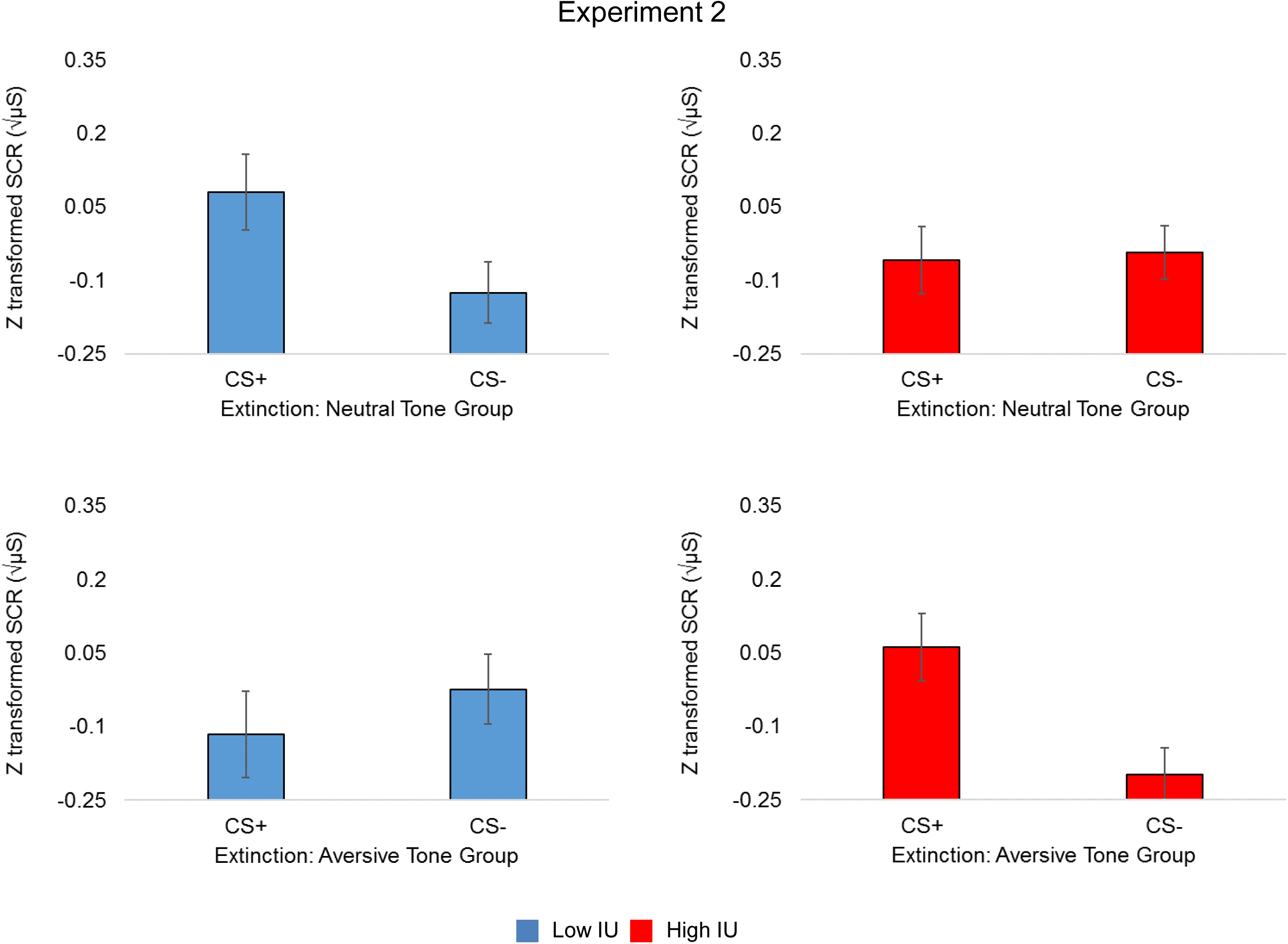
Bar graphs depicting IU estimated at + or - 1 SD of mean IU (controlling for STAI) from the multilevel model analysis for SCR magnitude during extinction. In experiment 2, high IU, relative to low IU individuals were only found to show larger skin conductance responding to the learned cues with threatening properties during extinction i.e. mildly aversive tone. Bars represent standard error at + or – 1 SD of mean IU. Square root transformed and z-scored SCR magnitude (μS), skin conductance magnitude measured in microSiemens

Moreover, low IU individuals in the aversive tone group had larger SCR magnitude in early (*M* = .08, *SE* = .07) vs. late extinction (*M* = -.21, *SE* = .08), *p* = .010, as well as reduced SCR magnitude in late extinction (*M* = -.21, *SE* = .08), compared to low IU individuals in the neutral tone group (*M* = .3, *SE* = .07), *p* =.42 [Group × Time × IU, *F*(1, 287.741) = 4.530. *p* = .034]. No other significant main effects of Group or interactions with IU or STAI emerged, max *F* = 3.391.

## Experiment 2: Conclusion

For experiment 2 we observed typical profiles of acquisition and extinction, where larger SCR magnitudes and expectancy ratings were found for the CS+ vs. CS-, despite the different threat levels of the US.

High IU was only associated with larger SCR magnitude to the CS+ versus CS-with threatening properties during extinction i.e. the mildly aversive tone. Conversely, low IU individuals in the neutral tone group displayed larger SCR magnitude to the CS+ versus CS-during extinction.

Similar to experiment 1, low STAI was associated with greater discrimination of expectancy of the tones with the CS+ vs. CS-, compared to high STAI. This effect occurred regardless of tone group.

## General Discussion

In the current research, we show that differences in self-reported IU are related to extinction depending on the level of uncertain threat present. These results partially replicate and extend prior findings from our lab of bodily and neural responding associated with IU and threat extinction (Morriss, Christakou, & van Reekum, 2015, 2016; Morriss, Macdonald, & van Reekum, 2016). Importantly, these findings provide another piece of the puzzle in recognising the relevance of IU-related mechanisms in disrupting threat extinction, which will likely have implications for current and future anxiety disorder diagnosis and treatment targets.

For both experiments we observed typical patterns of acquisition and extinction, where larger SCR magnitudes and expectancy ratings were found for the CS+ vs. CS-. In the first experiment, we aimed to examine the effect of an aversive uncertain US (i.e. human scream) on threat extinction and individual differences in IU. The aversive US was presented with a 50% reinforcement schedule during acquisition. High IU was associated with larger SCR magnitude to CS+ versus CS-cues during extinction. This finding is line with previous research examining IU and threat extinction (Dunsmoor, Campese, Ceceli, LeDoux, & Phelps, 2015; Lucas, Luck, & Lipp, 2018; Morriss, Christakou, & van Reekum, 2015, 2016; Morriss, Macdonald, & van Reekum, 2016). In the second experiment, we kept the same partial reinforcement procedure but changed the aversiveness of the US: One group of participants received a benign tone and another group of participants received a mildy aversive tone. In this experiment, high IU was only associated with larger SCR magnitude to the learned cues with threatening properties (i.e. the mildly aversive tone) during extinction. Interestingly, individuals reporting *low* IU in the neutral tone group displayed larger SCR magnitude to the CS+ versus CS-during extinction. This result was unexpected and warrants further investigation. The observed IU-related effects on SCR magnitude during extinction for experiment one and two were specific to IU, over STAI.

Taken together, the results from experiment one and two suggest that uncertain threat, even when it is mild, is an important factor in disrupting extinction in high IU individuals, as indexed by skin conductance. From a clinical perspective, these findings are particularly interesting, as associative learning principles underlie exposure-based therapies. For example, we can speculate that patients undergoing exposure therapy may require a different number of sessions depending on their IU score and the perceived aversiveness of the conditioned stressor(s).

In the current experiments we did not observe time-based effects of IU and threat extinction (Morriss, Christakou, & van Reekum, 2015, 2016). The difference between these experimental findings may be due to the reinforcement rate and timing of the CS. In the current experiments, we used a 50% reinforcement rate during the acquisition phase, whilst in our original experiments the rate was 100%. We used a 50% reinforcement rate in order to assess the conditioned response without the confound of the sound and to maintain the effect of conditioning longer during extinction, which, we suggest, is maintained particularly for high IU individuals. In addition, the current experiments used a CS of 4 seconds, whilst in our original experiments the CS was 1.5 seconds. It is advantageous to use a CS with a longer duration as it allows for more SCRs to be captured across all trials. Despite these design differences, IU-related effects were still observed in extinction.

Interestingly, our IU-related results differed depending on the type of measurement we used. The IU-related results in extinction were consistent for SCR magnitude across experiments one and two. The majority of research examining the effects of IU on threat acquisition and extinction have found significant relationships between IU and psychophysiological measures such as startle and skin conductance (Chin, Nelson, Jackson, & Hajcak, 2016; Morriss, Christakou, & van Reekum, 2015, 2016; Morriss, Macdonald, & van Reekum, 2016; Morriss, McSorley, & van Reekum, 2017; Sjouwerman, Scharfenort, & Lonsdorf, 2017). For the ratings we observed results with STAI, over IU, in experiment one and two. In experiment one, for both acquisition and extinction, individuals scoring higher on trait anxiety tended to have higher ratings of expectancy for both the CS+ and CS-, whilst individuals lower on trait anxiety showed greater discrimination between expectancy of the CS+ versus CS-. In experiment two, STAI significantly predicted the expectancy ratings during extinction. To our knowledge only a few studies have observed IU effects on ratings during acquisition and extinction (Morriss, Macdonald, & van Reekum, 2016; Sjouwerman, Scharfenort, & Lonsdorf, 2017). We therefore think that IU may be a more suitable predictor of bodily responses during threat extinction. The lack of consistent patterns between psychophysiological and rating measures for IU may, at least in our studies, also be due to the time between phasic cue events and rating periods in the experiment, where recall of expectancy was required for each block at the moment of rating.

A few issues with the current experiments should be further addressed in future research to assess the robustness and generalizability of the findings reported here. Firstly, the generality of these findings should be tested in future studies using stimuli that vary in aversiveness other than sounds e.g. level of shock, fearful/angry faces. Secondly, the sample contains mainly young female participants, and future studies should look to test more diverse samples.

In conclusion, these initial results provide some insight into how threat level and extinction may be related to IU, which may be relevant for understanding uncertainty-induced anxiety diagnostics and treatment targets (Carleton, 2016a, 2016b; Grupe & Nitschke, 2013). Further research is needed to explore how individual differences in IU modulate learned associations that vary in valence and arousal.

## Footnotes

1 The effect of Stimulus × STAI during acquisition was only observed when IU was included in the model.

## References

Ben-Shakhar, G. (1985). Standardization within individuals: A simple method to neutralize individual differences in skin conductance. Psychophysiology, 22(3), 292–299.

Carleton, R. N. (2016a). Fear of the unknown: One fear to rule them all? Journal of Anxiety Disorders, 41, 5–21.

Carleton, R. N. (2016b). Into the unknown: A review and synthesis of contemporary models involving uncertainty. Journal of Anxiety Disorders, 39, 30–43.

Chin, B., Nelson, B. D., Jackson, F., & Hajcak, G. (2016). Intolerance of uncertainty and startle potentiation in relation to different threat reinforcement rates. International Journal of Psychophysiology, 99, 79–84.

Craske, M. G., Treanor, M., Conway, C. C., Zbozinek, T., & Vervliet, B. (2014). Maximizing exposure therapy: an inhibitory learning approach. Behaviour Research and Therapy, 58, 10–23.

Dawson, M. E., Schell, A. M., & Filion, D. L. (2000). The Electrodermal System. In J. T. Cacioppo, L. G. Tassinary, & G. G. Berntson (Eds.), Handbook of Physiology (2nd ed., pp. 200–223). Cambridge, UK: Cambridge University Press.

Duits, P., Cath, D. C., Lissek, S., Hox, J. J., Hamm, A. O., Engelhard, I. M.,…Baas, J. M. (2015). Updated meta-analysis of classical fear conditioning in the anxiety disorders. Depression and Anxiety, 32(4), 239–253.

Dunsmoor, J. E., Campese, V. D., Ceceli, A. O., LeDoux, J. E., & Phelps, E. A. (2015). Novelty-facilitated extinction: providing a novel outcome in place of an expected threat diminishes recovery of defensive responses. Biological Psychiatry, 78(3), 203–209.

Etkin, A., & Wager, T. D. (2007). Functional neuroimaging of anxiety: a meta-analysis of emotional processing in PTSD, social anxiety disorder, and specific phobia. The American Journal of Psychiatry, 164(10), 1476.

Freeston, M. H., Rhéaume, J., Letarte, H., Dugas, M. J., & Ladouceur, R. (1994). Why do people worry? Personality and Individual Differences, 17(6), 791–802.

Grupe, D. W., & Nitschke, J. B. (2013). Uncertainty and anticipation in anxiety: an integrated neurobiological and psychological perspective. Nature Reviews Neuroscience, 14(7), 488–501.

LeDoux, J., & Daw, N. D. (2018). Surviving threats: neural circuit and computational implications of a new taxonomy of defensive behaviour. Nature Reviews Neuroscience.

Leonard, D. W. (1975). Partial reinforcement effects in classical aversive conditioning in rabbits and human beings. Journal of Comparative and Physiological Psychology, 88(2), 596.

Li, Y., Nakae, K., Ishii, S., & Naoki, H. (2016). Uncertainty-dependent extinction of fear memory in an amygdala-mPFC neural circuit model. PLoS computational biology, 12(9), e1005099.

Livneh, U., & Paz, R. (2012). Amygdala-prefrontal synchronization underlies resistance to extinction of aversive memories. Neuron, 75(1), 133–142.

Lonsdorf, T. B., Menz, M. M., Andreatta, M., Fullana, M. A., Golkar, A., Haaker, J.,… Kruse, O. (2017). Don’t fear ‘fear conditioning‘: Methodological considerations for the design and analysis of studies on human fear acquisition, extinction, and return of fear. Neuroscience & Biobehavioral Reviews.

Lonsdorf, T. B., & Merz, C. J. (2017). More than just noise: Inter-individual differences in fear acquisition, extinction and return of fear in humans-Biological, experiential, temperamental factors, and methodological pitfalls. Neuroscience & Biobehavioral Reviews, 80, 703–728.

Lucas, K., Luck, C. C., & Lipp, O. V. (2018). Novelty-facilitated extinction and the reinstatement of conditional human fear. Behaviour Research and Therapy, 109, 68–74.

Milad, M. R., & Quirk, G. J. (2012). Fear extinction as a model for translational neuroscience: ten years of progress. Annual review of psychology, 63, 129–151.

Morriss, J., Chapman, C., Tomlinson, S., & Van Reekum, C. M. (2018). Escape the bear and fall to the lion: The impact of avoidance availability on threat acquisition and extinction. Biological Psychology, 138, 73–80.

Morriss, J., Christakou, A., & Van Reekum, C. M. (2015). Intolerance of uncertainty predicts fear extinction in amygdala-ventromedial prefrontal cortical circuitry. Biology of Mood & Anxiety Disorders, 5(1), 1.

Morriss, J., Christakou, A., & Van Reekum, C. M. (2016). Nothing is safe: Intolerance of uncertainty is associated with compromised fear extinction learning. Biological Psychology, 121, 187–193.

Morriss, J., Macdonald, B., & van Reekum, C. M. (2016). What Is Going On Around Here? Intolerance of Uncertainty Predicts Threat Generalization. PloS one, 11(5), e0154494.

Morriss, J., McSorley, E., & van Reekum, C. M. (2017). I don‘t know where to look: the impact of intolerance of uncertainty on saccades towards non-predictive emotional face distractors. Cognition and Emotion, 1–10.

Neumann, D. L., Waters, A. M., & Westbury, H. R. (2008). The use of an unpleasant sound as the unconditional stimulus in aversive Pavlovian conditioning experiments that involve children and adolescent participants. Behavior Research Methods, 40(2), 622–625.

Phelps, E. A., Delgado, M. R., Nearing, K. I., & LeDoux, J. E. (2004). Extinction learning in humans: role of the amygdala and vmPFC. Neuron, 43(6), 897–905.

Sjouwerman, R., Scharfenort, R., & Lonsdorf, T. B. (2017). Individual differences in fear learning: Specificity to trait-anxiety beyond other measures of negative affect, and mediation via amygdala activation. bioRxiv, 233528.

Spielberger, C. D., Gorsuch, R. L., Lushene, R., Vagg, P., & Jacobs, G. (1983). Consulting Psychologists Press, Inc. 2». Palo Alto (CA).

